# Passive copper release from ornamental rain chains is harmful to mosquito larvae and deters egg-laying

**DOI:** 10.64898/2026.07.26.740750

**Authors:** Claudio R. Lazzari, Bertrand Le Mellat

## Abstract

Urban mosquitoes use accumulated rainwater in private gardens and public green spaces for reproduction. We found that ornamental copper chains, which direct rainwater from gutters into open containers, can significantly reduce the number of mosquito larvae in the collected water, sometimes eliminating them entirely. Although the impact of copper salts or copper objects in water on mosquito larvae is well documented, a detailed analysis of the effects of dissolved copper in terms of concentration and biological impact has yet to be conducted. In this study, we present the results of two experimental analyses: one conducted in gardens and the other in the laboratory. These experiments were designed to corroborate consistent, albeit circumstantial, observations, and to describe the sensitivity of mosquito larvae to rainwater containing low quantities of copper. Field experiments were conducted during a period of high tiger mosquito (*Aedes albopictus*) abundance, while laboratory experiments used the yellow fever mosquito (*Aedes aegypti*) as a biological model. Our field experiments revealed that rain chains release sufficient copper into the water to drastically reduce the number of juvenile mosquitoes. In the laboratory, rainwater containing copper released by the chains affected hatching and larval development; however, the most significant effect was that females laid almost no eggs in the presence of copper-containing water.

## 1. Introduction

Urban mosquitoes reproduce in standing outdoor water. Even small artificial or natural receptacles constitute adequate places for females to lay their eggs and for larval development. Two cosmopolitan and invasive species that are well adapted to urban environments are the Asian tiger mosquito (*Aedes albopictus*) and the yellow fever mosquito (*Aedes aegypti*). These species are major vectors of several pathogens, including dengue, chikungunya, Zika and yellow fever, as well as other arboviruses that infect humans. They also carry *Dirofilaria* worms that infect domestic animals.

Although avoiding the accumulation of water outdoors is an effective way to control the proliferation of mosquitoes in gardens and peri-domestic areas, this is not always simple or desirable. Rainwater is frequently collected in gardens for decorative or practical purposes, such as watering plants and the containers also facilitate mosquito proliferation.

The detrimental effects of copper salts and metallic copper on mosquito larvae are well documented in the literature. Several studies analysing the biological effects of dissolved metallic copper on mosquito larval development have been published [1– 6]. Negative impacts on larval development and increased mortality have been observed at relatively low concentrations of the metal in both field studies and controlled laboratory experiments.

One of the authors of this study (B. L. M.), who sells ornamental rain chains, received feedback from customers who had installed copper chains in their gardens. These clients reported that adult mosquitoes appeared to rarely approach containers collecting rainwater that had flowed through the chains and that fewer larvae and pupae seemed to develop in this water than in containers collecting rainwater directly without a chain. While these observations were consistent with published scientific evidence, they raised questions about the biological significance and underlying mechanisms of the observed effect of rain chains.

This study aimed to shed additional light on the potential effectiveness of copper rain chains in reducing the density of mosquitoes in domestic and peridomestic environments. Specifically, the study sought to: Firstly, we evaluated any potential adverse impacts at various stages, including egg hatching, larval development, and egg-laying. Secondly, we aimed to quantify the amount of copper accumulated in water collected from roofs and flowing through rain chains.

## 2. Materials and Methods

### 2.1. Field experiments

The company *Chaînes de Pluie*, based in Le Mans, France, provided different models of decorative rain chains made of copper. Figure 1 shows an example of a chain and how it is installed to conduct rainwater collected by gutters from a roof.

**Figure 1.**
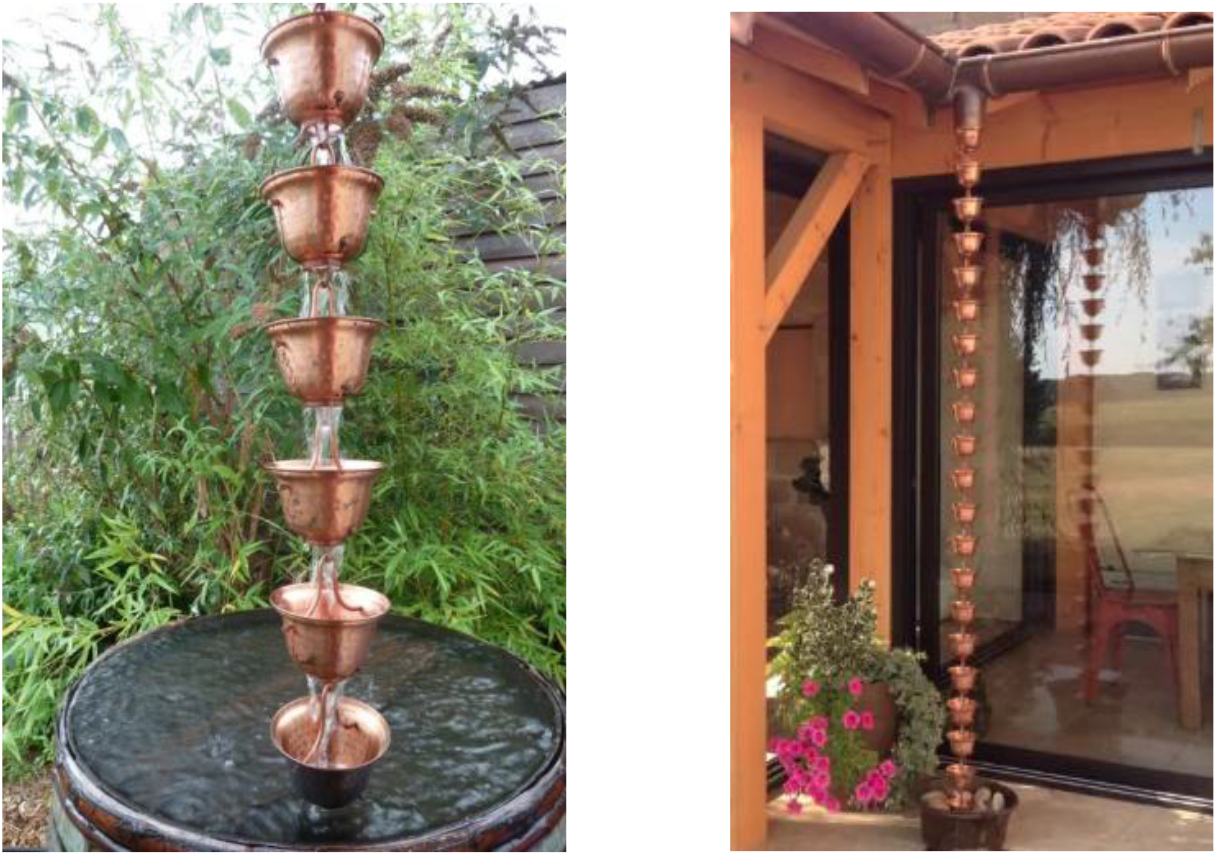
The left picture depicts the “Shinjuku” chain model, and right one illustrates how decorative rain chains are installed to collect rainwater from roof gutters and direct it into barrels or other types of rainwater collector.

Further information can be found on the company’s website: https://www.chainesdepluie.com/.

Chains were installed on gutters to collect rainwater from roofs and direct it into containers located in private residences and garden sheds in the French departments of Indre-et-Loire and Sarthe. At each site, two identical containers were placed side by side to collect rainwater flowing through the chains or falling directly. A total of 18 experiments were conducted between May and September 2025, during a period of high tiger mosquito abundance in France.

Water was allowed to accumulate for varying periods according to the rainfall pattern, giving mosquitoes time to lay eggs and allowing the larvae to hatch and grow. This process takes roughly 10 to 14 days after a rainfall event. The containers were then carefully inspected for immature mosquito stages and the copper content of the water was measured using a *JBL Pro Aquatest Cu* kit (Germany).

Since the installations and follow-ups were conducted in different locations by different observers, and since the water was collected in different types of containers, two categories of data were established. The integrity of the observation points was considered in order to conduct a qualitative assessment of mosquito behaviour and ensure consistency of observation. For quantitative and statistical analyses, a subset of sites using standardised black containers with a volume of 3.5 litres (usually employed in mosquito oviposition traps) located in Indre-et-Loire (47°20’42”N – 0°39’16”E) and observed by the same researcher (C. L.) were used. As data were collected throughout the season involving two types of structures and three chain models, a pairwise statistical analysis was performed to compare the presence of mosquitoes in pairs of control and experimental containers tested simultaneously.

**Table 1.** Observation sites, observers, and conditions of outdoor essays on the presence of mosquitoes in water flowing through copper rain chains. The column labelled “Area” indicates the surface area of the structure collecting rainwater.

| Responsible | Region | Chain model | Structure | Area | Period |
| --- | --- | --- | --- | --- | --- |
| Collaborator #1 | Marne-et-Loire | Ôtemachi | Shelter | 10m<sup>2</sup> | Spring 2024 |
| Collaborator #2 | Sarthe | Sinjuku | Roofed yard | 20m<sup>2</sup> | Spring 2025 |
| Collaborator #3 | Sarthe | Kinkakuji | Barn | 15m<sup>2</sup> | Spring 2025 |
| Collaborator #4 | Sarthe | Shinjuku | Veranda | 12m<sup>2</sup> | Spring 2025 |
| Collaborator #5 | Sarthe | Shinjuku | House | 50m<sup>2</sup> | Spring 2025 |
| Collaborator #6 | Sarthe | Ôtemachi | House | 50m<sup>2</sup> | Summer 2025 |
| B. Le Mellat | Sarthe | Ryogoku | Shelter | 12m<sup>2</sup> | Spring 2025 |
| B. Le Mellat | Sarthe | Ryogoku | Shelter | 12m<sup>2</sup> | Summer 2025 |
| C. Lazzari | Indre-et-Loire | Shinjuku | House | 40m<sup>2</sup> | Summer 2025 |
| C. Lazzari | Indre-et-Loire | Shijuku | House | 40m<sup>2</sup> | Summer 2025 |
| C. Lazzari | Indre-et-Loire | Oji | Shelter | 10m<sup>2</sup> | Summer 2025 |
| C. Lazzari | Indre-et-Loire | Ôtemachi | Shelter | 10m<sup>2</sup> | Summer 2025 |
| C. Lazzari | Indre-et-Loire | Shinjuku | House | 40m<sup>2</sup> | Summer 2025 |
| C. Lazzari | Indre-et-Loire | Oji | Shelter | 10m<sup>2</sup> | Summer 2025 |
| C. Lazzari | Indre-et-Loire | Ôtemachi | Shelter | 10m<sup>2</sup> | Summer 2025 |
| C. Lazzari | Indre-et-Loire | Shinjuku | House | 40m<sup>2</sup> | Summer 2025 |
| C. Lazzari | Indre-et-Loire | Oji | Shelter | 10m<sup>2</sup> | Summer 2025 |
| C. Lazzari | Indre-et-Loire | Ôtemachi | Shelter | 10m<sup>2</sup> | Summer 2025 |

### 3.2. Laboratory experiments

These studies were conducted under controlled conditions at the *Institut de Recherche sur la Biologie de l’Insecte* in Tours, France.

*Aedes aegypti* belonging to the Bora strain and sensitive to insecticides were reared in the laboratory and kept under controlled conditions of temperature (26 ± 1 °C), relative humidity (60–70%) and illumination (12 hours of light and 12 hours of darkness).

The eggs were kept dry and placed in dechlorinated tap water until they hatched. The larvae were fed dry tropical fish food. The adults were kept in cages with free access to a 10% sugar solution in water. The females were fed an artificial blood meal replacement, *SkitoSnack* (Gonzalez et al., 2018; Kandel et al., 2020), which was prepared in our laboratory, and offered using an artificial feeder (*Hemotek*, UK). Our laboratory adopted this highly efficient method several years ago, resulting in satisfactory survival and egg production rates for several generations.

To ensure consistency across the different experiments, rainwater was placed in one-litre containers and sections of copper chain were submerged for four weeks. After this time, the copper concentration reached 1.2 mg/L. No copper was detected in the control sample.

Groups of eggs were placed in small plastic containers. One group contained 10 ml of rainwater that had been in contact with copper rain chains, while the other group contained rainwater that had been collected directly. Different experimental series were conducted, with some larvae receiving food and others not. Hatching, survival and development were then evaluated simultaneously in an equal number of containers from both the experimental and control groups.

The oviposition preference of the females was evaluated as follows: Groups of around 100 females were fed artificial blood and offered recipients containing a 5 cm wide, 15 cm long piece of filter paper soaked in either rainwater containing 1.2 mg/L of copper (Cu) or rainwater without Cu (the control group) for egg-laying over the next four days. Four repetitions of this experiment were conducted.

## 3. Results

### 3.1. Field experiments

A noticeable reduction in numbers was observed, even total suppression, of mosquito larvae and pupae in the water, and in the number of adult mosquitoes flying close to the surface, in containers collecting rainwater flowing through rain chains, compared to containers collecting rainwater directly. These effects of the presence of rain chains were consistently reported by independent observers at all experimental sites.

The amount of copper that accumulated varied according to the rainfall pattern and chain model at each observation site. We obtained values ranging from 0.6 to 1.6 mg/L. The most frequent values were between 0.8 and 1.2 mg/L.

As outlined in the Materials and Methods section, it was possible to make precise comparisons based on a subset of ten well-controlled essays. Figure 2 depicts the results, which reveal statistically significant differences in the pairwise comparisons between the control and experimental conditions.

**Figure 2.**
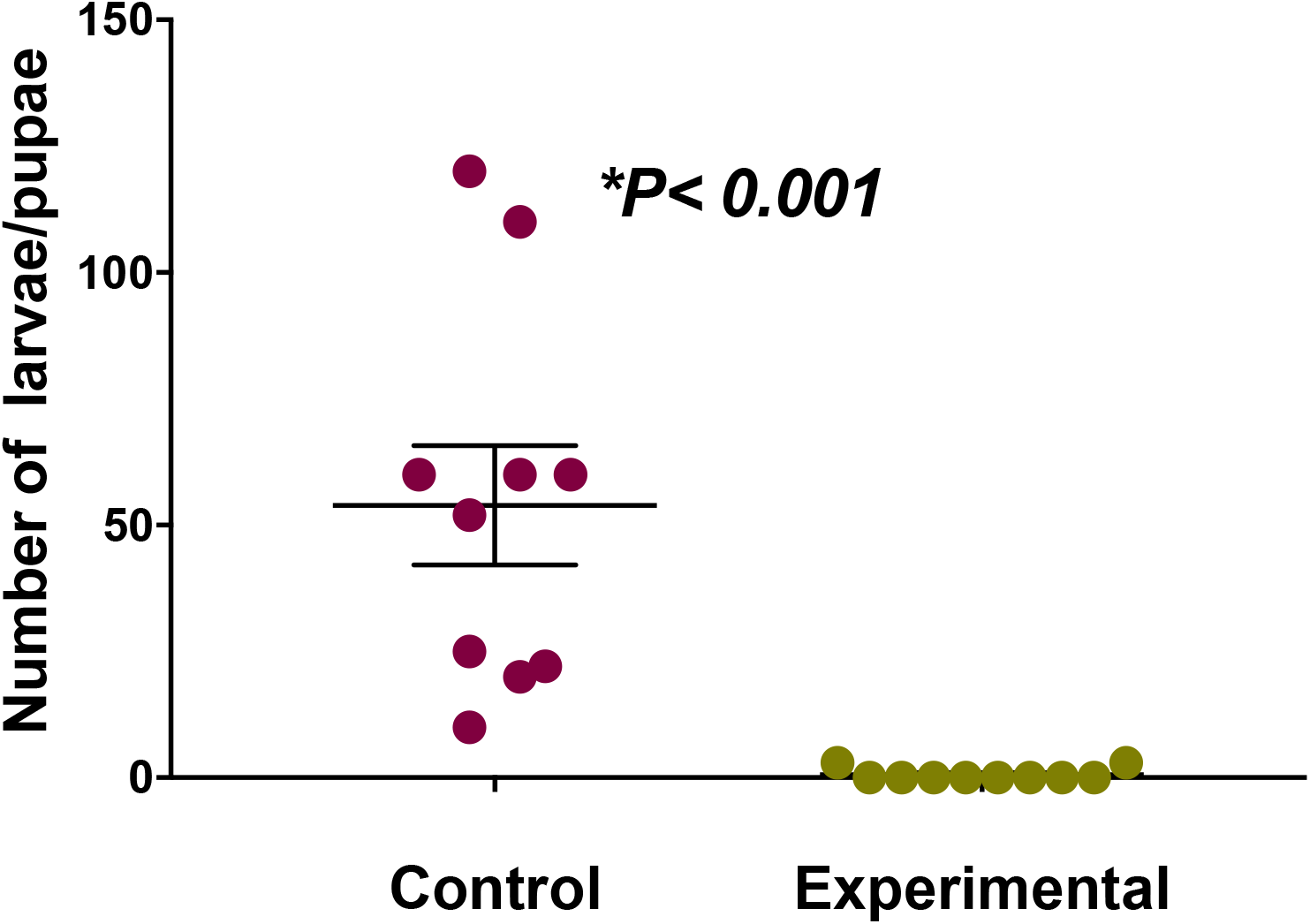
Comparison in the number of larvae and pupae in the control container, filled with rainwater collected directly and an identical container, filled with rainwater flowing along a copper rain chain. Recipients were placed side-by-side and a paired *t*-test was performed to compare both situations. The graph depicts individual data, the mean ± s.e.m of each condition (n= 10).

Video S1 shows a recurring observation at the different sites: more females were found flying close to the water surface in the control containers than in the experimental ones.

### 3.2. Laboratory experiments

#### 3.2.1. Egg hatching and larval development

The presence of copper in the water was found to have an observable impact on egg hatching. Although hatching occurred in both treatments, the number of larvae in Cu-loaded rainwater was lower. However, this difference was not statistically significant and could not explain by itself the near-complete absence of mosquito larvae in chain-collected rainwater in our outdoor experiments.

Larvae in both groups survived when food was added to the water (n = 40). However, while most of the control group reached the fourth instar within a week, those exposed to copper-containing rainwater only reached the second instar after two weeks, indicating a significant growth delay. Even though in line with findings reported by previous authors, this effect alone does not explain the virtual absence of mosquito larvae in rainwater containing copper observed in the field.

To test whether the reduction in mosquito larvae could be due to microorganisms in the water having a negative impact on them, we ran an experiment in which the larvae were not fed. When no additional food was added to the water (n = 10), a similar result to that obtained previously was verified. Larval development was retarded in both treatments, and not significant differences were observed.

#### 3.2.2. Egg-laying

The most notable difference between the treatments was related to egg laying. While *Aedes aegypti* females laid a high number of eggs on oviposition papers impregnated with control rainwater, very few or no eggs were deposited on papers soaked in rainwater containing copper, as shown in Figure 3.

**Figure 3.**
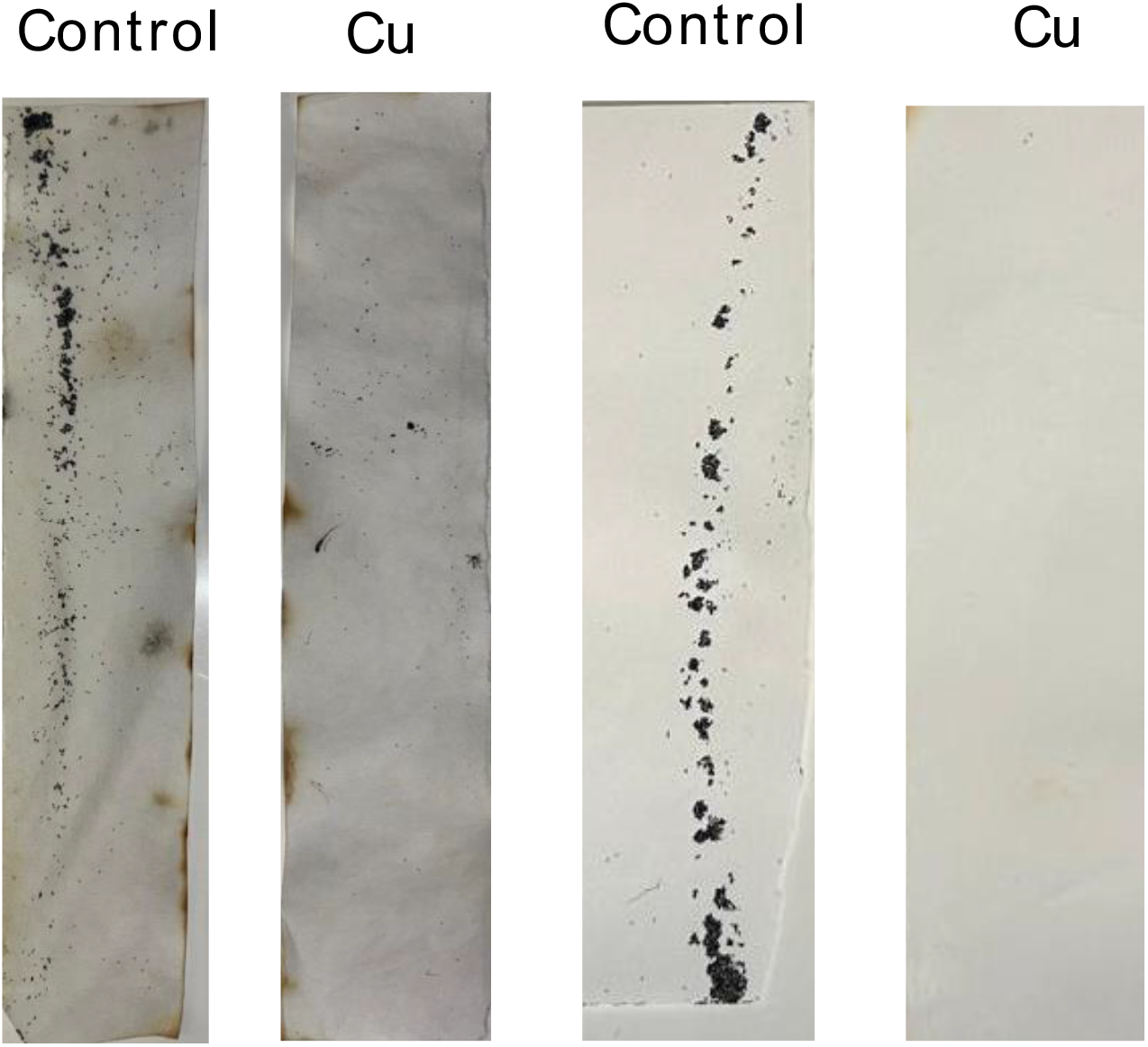
Oviposition papers were soaked in rainwater, either untreated (control) or containing 1.2 mg/L of copper. Two sample records are presented.

## 4. Discussion

The experiments presented here demonstrate that rainwater flowing along copper chains can leach enough metal to render the water unsuitable for colonisation by at least two mosquito species of major epidemiological importance: *Aedes albopictus* (tiger mosquito) and *Aedes aegypti* (yellow fever mosquito).

Previous studies have documented the toxic effects of dissolved copper in aquatic environments, whether introduced as salts or in metallic form [1–6]. These properties have been successfully exploited for mosquito control in settings characterized by multiple breeding sites, such as cemeteries [3–6]. Given its capacity to impair mosquito development, copper represents a promising tool for reducing mosquito abundance in urban environments.

Our findings extend this body of knowledge by showing that ornamental rain chains can act as passive copper delivery systems, limiting colonisation of both decorative and functional water containers. Importantly, our study also provides insight into the mechanisms underlying this effect.

Consistent with earlier reports, we observed reduced egg hatching, decreased larval survival, and delayed development in copper-exposed conditions [4-5]. However, while these effects have been widely reported, they do not fully account for the near absence of larvae observed in our outdoor experiments with copper-treated water.

Notably, our results indicate that the most pronounced effect of copper was the inhibition of oviposition by gravid females, a phenomenon that has not been previously described. This suggests that copper may act not only as a toxicant but also as a deterrent influencing habitat selection.

One possible explanation for this behavioural avoidance lies in the well-documented antimicrobial properties of metallic copper [7]. Oviposition site selection in mosquitoes is influenced by a combination of abiotic and biotic factors, including water chemistry and container characteristics [8]. In particular, microbial communities within water-filled habitats play a critical role in shaping oviposition preferences in Aedes aegypti. Volatile compounds produced by these microorganisms can act as attractants, guiding females to suitable egg-laying sites [9–12]. Additionally, bacteria and their metabolic byproducts can alter physicochemical properties of the water, such as dissolved oxygen levels, and have been shown to promote egg hatching [13].

Conversely, gravid females themselves contribute to shaping the microbial composition of breeding sites during oviposition, introducing microorganisms that may enhance larval development [14–15]. The presence of copper, even at low concentrations, could disrupt these microbial communities, thereby reducing both the attractiveness of the site and its suitability for offspring development. This dual action of direct toxicity and indirect interference with microbial cues, may explain the strong avoidance behaviour observed in our study.

## 5. Conclusions

In conclusion, rainwater passing over copper chains can accumulate sufficient dissolved copper to significantly disrupt the colonisation of artificial water containers by *Aedes albopictus* and *Aedes aegypti*. Beyond confirming previously reported toxic effects on immature stages, our findings reveal that copper also exerts a strong deterrent effect on oviposition, substantially reducing the likelihood that gravid females will select treated water as a breeding site.

This dual mode of action, combining direct toxicity with behavioural avoidance, provides a compelling explanation for the near absence of larvae observed under field conditions. By likely altering microbial communities and the associated chemical cues that guide oviposition, copper-treated water becomes both inhospitable and unattractive to mosquitoes. These results highlight the importance of considering not only lethal effects but also sublethal and ecological mechanisms when evaluating vector control strategies.

From an applied perspective, the use of copper-based materials such as ornamental rain chains represents a simple, passive, and potentially cost-effective intervention for reducing mosquito breeding in urban environments. Such approaches could be particularly valuable in settings where numerous small water-holding containers are difficult to manage through conventional control methods. However, further research is needed to determine the long-term efficacy, optimal deployment conditions, and potential environmental impacts of copper release in diverse urban contexts.

Overall, our study supports the integration of copper-mediated strategies into broader mosquito management programs and underscores the value of leveraging material properties and ecological interactions to develop innovative and sustainable vector control solutions.

## Supporting information

Adults flying over the surface of treated and untreated rainwater.

## Supplementary Materials

Video S1: adults flying over the surface of treated and untreated rainwater.

## Author Contributions

Conceptualization, C.L. and BLM.; methodology, C.L.; validation, C.L.; formal analysis, C.L.; investigation, C.L.; resources, C.L. and B.L.M.; data curation, C.L.; writing—original draft preparation, C.L.; writing—review and editing, C.L. and B.L.M.; visualization, C.L. and B.L.M.; supervision, C.L.; project administration, C.L. and B.L.M.; funding acquisition, C.L. and B.L.M. Both authors have read and agreed to the published version of the manuscript

## Acknowledgments

The authors express their gratitude to Esteban Moyer for his help in maintaining *Aedes aegypti* in the lab, as well as to Mr Stéphane Marie, Ms Pangole, Mr and Ms Rivalain, Mr and Ms Guittière, Mr and Ms Carpentier, Ms Renard and Mr and Ms Berrou, and Dr. T. Insausti, for accepting the presence of chains and water recipients at their particular gardens. David Carrasco and the technical personnel of MIVEGEC (IRD-CNRS, Montpellier, France) generously provided eggs of *Aedes aegypti* used in the experiments. *Le Mans Innovation* has provided its support for obtaining funds. The IRBI (CNRS-Univ. Tours) provided the facilities for the insectarium and experimental rooms. The REPEL project, financed by INEE-CNRS partially contributed to the maintenance of mosquitoes in the laboratory.

